# Massive Multiplexing Can Deliver a $1 Test for COVID-19

**DOI:** 10.1101/2020.05.05.079400

**Authors:** Paul DN Hebert, Sean WJ Prosser, Natalia V Ivanova, Evgeny V Zakharov, Sujeevan Ratnasingham

**Affiliations:** Centre for Biodiversity Genomics, University of Guelph, 50 Stone Road, Guelph, ON, N1G 2W1; Department of Integrative Biology, University of Guelph, 50 Stone Road, Guelph, ON, N1G 2W1

**Keywords:** Sequel, ION S5, E gene, SARS-CoV-2, coronavirus, screening, group tests

## Abstract

The severe acute respiratory syndrome virus, SARS-CoV-2 (hereafter COVID-19), rapidly achieved global pandemic status, provoking large-scale screening programs in many nations. Their activation makes it imperative to identify methods that can deliver a diagnostic result at low cost. This paper describes an approach which employs sequence variation in the gene coding for its envelope protein as the basis for a scalable, inexpensive test for COVID-19. It achieves this by coupling a simple RNA extraction protocol with low-volume RT-PCR, followed by E-Gel screening and sequencing on high-throughput platforms to analyze 10,000 samples in a run. Slight modifications to the protocol could support screening programs for other known viruses and for viral discovery. Just as the $1,000 genome is transforming medicine, a $1 diagnostic test for viral and bacterial pathogens would represent a major advance for public health.

## INTRODUCTION

The emergence of COVID-19 as a global pandemic (Helmy et al. 2020) has created the need for greatly expanded screening programs to aid contact tracing and to protect medical staff and other vulnerable groups (WHO 2020a). Rapid characterization of the COVID-19 genome (Wu et al. 2020) and the limited variation among isolates (Forster et al. 2020) enabled the prompt development of diagnostic tests. Most involve two phases; RT-qPCR is used for initial screening while sequencing is employed for confirmation (WHO 2020b). For example, one widely adopted protocol employs RT-qPCR to screen for a 112 bp amplicon of the E gene for initial diagnosis followed by Sanger sequencing of a 95 bp segment of the RdRp gene for confirmation (Corman et al. 2020, Public Health Ontario 2020). The adoption of standard protocols has aided quality assurance, but it has constrained the volume of testing because reagent kits are in short supply (Sharfstein et al. 2020) and costs ($50–$100 USD per test) are high (Medicare 2020). As well, because workflows are complex, analytical results can be delayed for a week, limiting their value for the suppression of community spread. Although cassette-based RT-qPCR assays can deliver results in less than an hour (Vashist 2020), these instruments process too few samples at a time (<10) to support the large-scale surveillance programs (i.e. 100,000 samples per day) planned by many nations (Department of Health & Social Care 2020). Until recently, the collection of samples was challenging as nasopharyngeal swabs require medical staff, but saliva is equally effective (To et al. 2020). As the sample acquisition bottleneck has been broken, there is a need to increase analytical capacity, and this will be achieved most easily by employing infrastructure and expertise in existing genomics facilities. A fully distributed solution involving many low-volume labs would bring complexity to quality assurance and control. System management would be greatly simplified if a few core facilities were established, each initially processing 10,000 samples per day with subsequent increase as need arises. To support this production strategy, analytical approaches must scale effectively. Solutions have been proposed, ranging from new technologies (Schmid-Burgk et al. 2020, Zhang et al. 2020) to augmented Sanger sequencing (Chandler-Brown et al. 2020), but they are not fully ready for implementation. By contrast, highly multiplexed amplicon sequencing has a well-proven capacity to characterize many thousands of samples in a run (Hebert et al. 2018).

Aside from having the capacity to meet production goals, screening protocols must be rapid, inexpensive, and reliable. Results should ideally be available within a day to curb community transmission and costs become increasingly important as surveillance programs expand. For example, a white paper from the Rockefeller Foundation (2020) proposes that the USA analyze 30 million individuals weekly at an estimated annual cost of $150 billion. Because reliability is also essential, protocols must minimize the incidence of false positives and negatives (Wickramaratna 2020, Winter 2020). False positives typically arise from cross contamination or when tests lack specificity while false negatives reflect a lack of sensitivity or analytical mishap. Achieving required sensitivity is complicated by the fact that viral loads vary greatly among individuals and throughout the course of infection (Lescure et al. 2020). For example, To et al. (2020) report million-fold variation (10^3^–10^9^ per ml) from symptomatic individuals. Presuming adequate sensitivity, false negatives can still arise because of analytical error, but this risk can be evaluated by including controls spiked with synthetic RNA. The risk of false positives can be evaluated by including blank controls, but the best defense involves running a separate assay on a fresh sample when the first test is positive.

The protocol reported here is based on the analysis of sequence variation in a target gene through gels and sequencing. Employing in-well controls and supported by informatics platforms that automate the interpretation of results and report assembly, a single facility can process 10,000 samples per day. The Results section begins by demonstrating that sequence variation in the E gene can underpin a reliable diagnostic test for COVID-19 and then describes cost-effective protocols to assess this variation.

## RESULTS

### SELECTION OF TARGET GENE

The 29,870 bp genome of COVID-19 codes for 12 genes (Wu et al 2020), five of which (E, N, Orf1ab, RdRp, S) have often been employed in diagnostic tests (Vashist 2020). The envelope gene (E) was selected for evaluation, an effort that began with an examination of sequence divergence patterns among all known lineages of *Betacoronavirus.* These taxa are assigned to five subgenera, but two (*Hibecovirus*, *Nobecovirus*) are only known from bats. The other three infect diverse mammals, and five of their component lineages cause respiratory disease in humans. These include COVID-19 and SARS in the subgenus *Sarbecovirus*, Human coronavirus HKU1 and Human coronavirus_OC43 in *Embecovirus*, and MERS in *Merbecovirus.* Examination of all E gene sequences for taxa of *Betacoronavirus* revealed considerable length variation (210– 270 bp), but enough conserved amino acids to allow reliable alignment. A NJ tree based on sequences for the E gene recovered each subgenus as monophyletic (Figure 1a) with deep divergences between taxa in different subgenera (mean = 49%; range = 26–56%). When comparison focused on members of a subgenus, divergences were less, but still much higher than intraspecific variation. For example, the two human coronaviruses (OC43, HKU1) of *Embecovirus* possess 35% divergence versus intraspecific divergences of 1.2% and 7.6% (the latter divergence reflects differences between strains of HKU1 from rodents and humans). Similarly, although SARS and COVID-19 are strains of a single species within *Sarbecovirus*, they have a mean divergence of 5.3% versus intrastrain values of less than 0.1% (Figure 1b). Reflecting this fact, there are 11 diagnostic substitutions and a consistent 3 bp deletion between most sequences of the E gene from COVID-19 and SARS. Examination of all genome sequences for COVID-19 in the GISAID database (Elbe and Buckland-Merrett 2017) on April 15, 2020 further revealed that 98.5% of the 6,300 most reliable genomes (high coverage, >29 kb) possessed an identical sequence for the E gene. Divergence among the remaining 1.5% was low with most SNPs only detected in a single sequence. Although a few SNPs were present in 2–7 sequences, they never reduced the number of diagnostic substitutions below 9 so the E gene was targeted for analysis.

**Figure 1:**
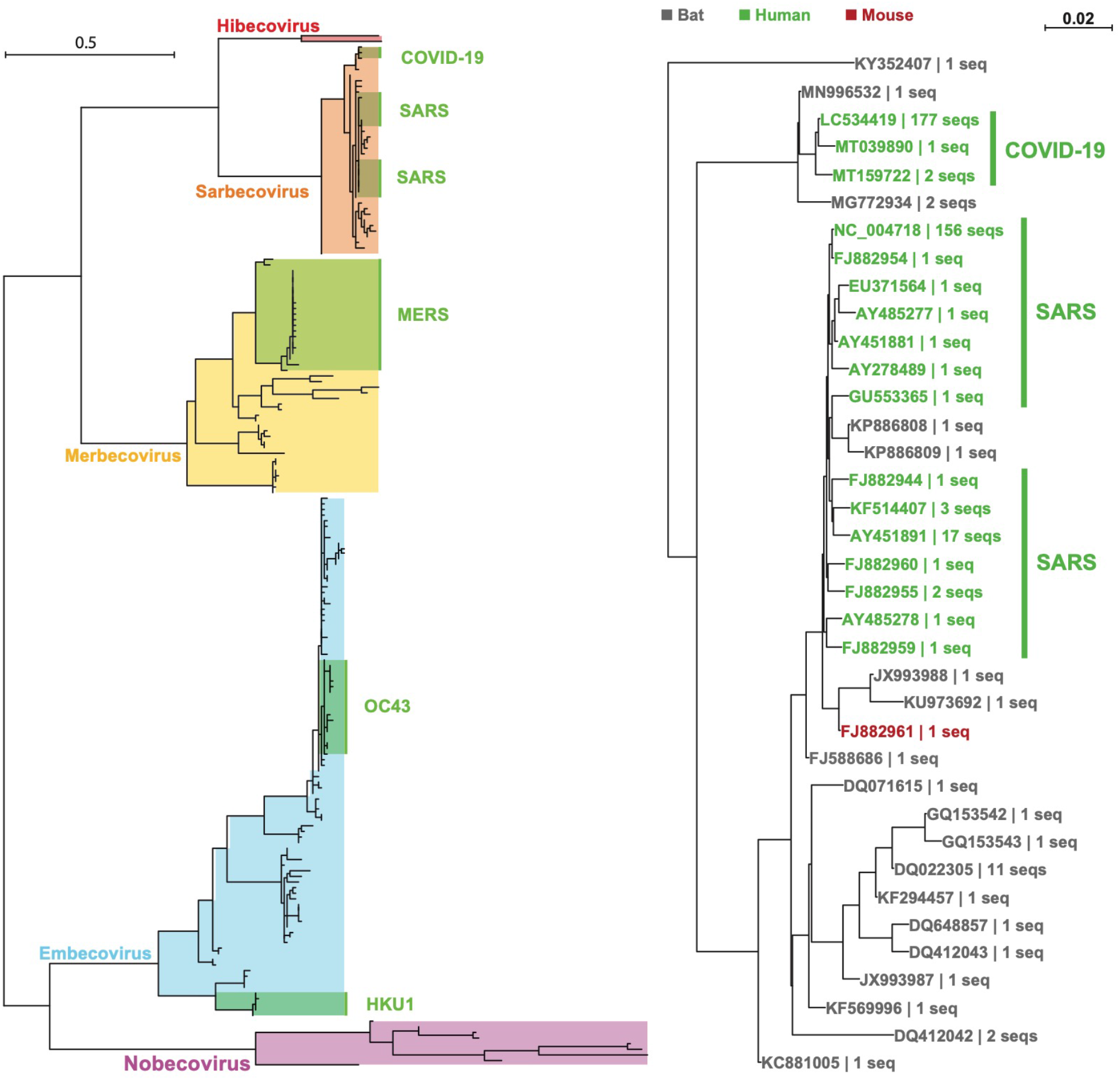
**A)** Neighbor-Joining (NJ) tree based on sequences for the E gene for all lineages of the genus *Betacoronavirus*. Lineages recovered from humans are highlighted in green. **B)** NJ tree based on all sequences of the E gene for members of the subgenus *Sarbecovirus*. Lineages from humans are in green while those only known from bats are in gray and the sole lineage from a mouse is in red.

### OVERVIEW OF PROTOCOL

The following text summarizes the six steps (Figure 2) in the screening protocol as well as material costs and required infrastructure. Next generation sequencing platforms (Sequel, Ion S5) are optimal for high production, but Sanger sequencing is cost-effective when fewer than 1,000 samples are tested per day. These workflows deliver a gel-based diagnostic within 8 hours and a sequence-based diagnostic within 24–48 hours depending on the platform employed.

**Figure 2:**
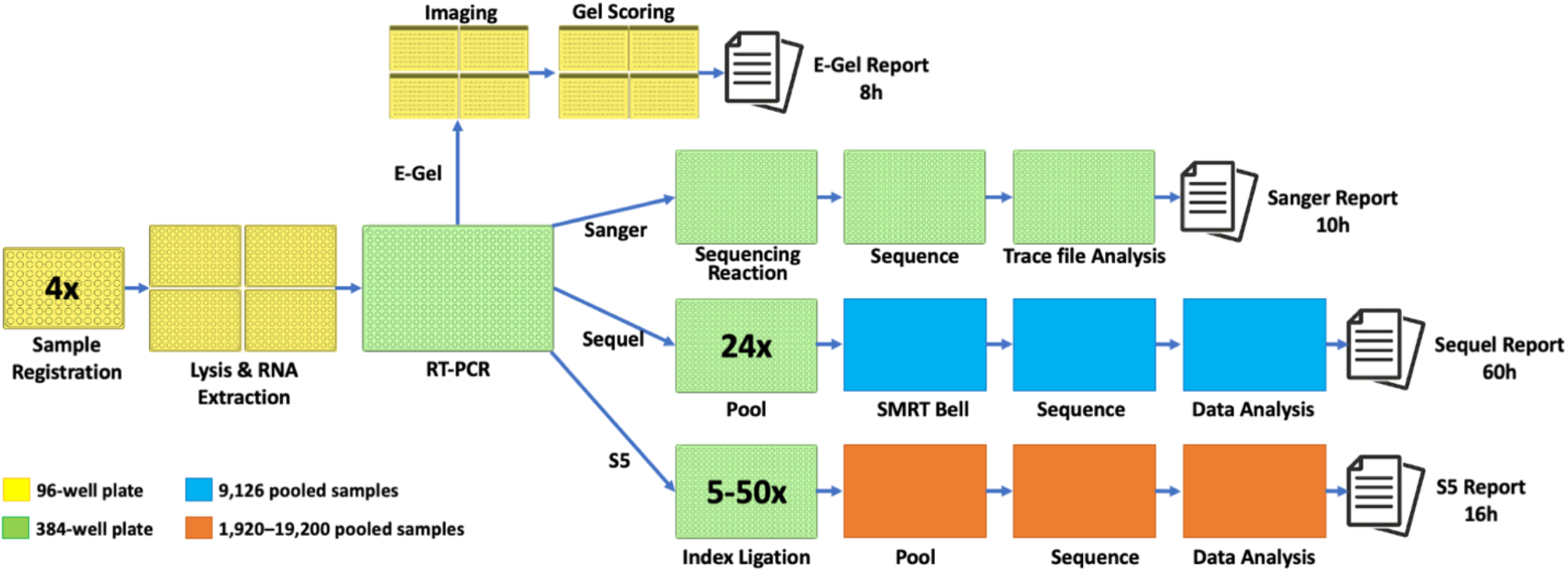
COVID-19 screening protocol. E-Gels are used to rapidly assess the presence of an amplicon for the E gene which is followed by sequence characterization. The three sequencing platforms have differing time requirements and production capacities.

#### 1. Sample Registration

Tubes containing universal viral medium or saliva are placed at 56°C for 30 minutes to inactivate the virus (Pastarino et al. 2020, Yang and Wang 2020). After heat treatment, tubes are centrifuged (2000 g for 1 minute) and then placed into 96-well racks where their 2-D barcodes are scanned to register each sample in a Laboratory Information Management System (LIMS). Each rack includes 94 test samples, 1 negative control, and one positive control.

Time to Complete: 1 h

Supplies: None

Infrastructure: 2 barcode scanners, LIMS

#### 2. Lysis and RNA Extraction

Step #2 is supported by a Biomek FX liquid handler whose deck is loaded with racked samples and plates (Extraction, Wash, Elution) with required consumables (magnetic beads, ethanol, wash buffers) as well as the RT-PCR plates required for Step #3. RNA extraction follows He et al. (2017) with minor modifications. Sample analysis begins with the transfer of 100 μl from each tube into the corresponding well of a 96-well Extraction plate (1.2 ml Abgene Square) pre-filled with 100 μl of guanidine thiocyanate buffer (4 M guanidinium thiocyanate, 160 mM dithiothreitol, 50 mM sodium citrate, 200 μg/ml of glycogen, 200 mM Tris-HCl, pH 8.9) held at 75°C on a Peltier block. After 15 min of lysis at this temperature, the plate is cooled to 20°C before 200 μl of 100% ethanol and 13 μl of magnetic beads (SpeedBeads, 10 mg/ml) are added to each well, at which point the FX mixes the beads and lysate by repeatedly (5x) aspirating/returning 90 μl aliquots from each well. The plate is then incubated on the deck at 20°C for 5 min to allow RNA/DNA to bind to the beads before they are harvested with a magnetic bead extractor (VP 407AM-N1-R-MagPin R). The extractor is covered with a disposable cover plate (VWR 82006-636) to allow bead recovery from multiple Extraction plates without contamination (Oberacker et al. 2019). A new cover plate is added before the extractor is submerged into each Abgene plate for 2 min to collect beads with bound DNA/RNA. Beads are then immersed in a Wash plate (Costar) pre-filled with 150 μl of 70% ethanol (in DEPC-treated water) by moving the extractor up and down three times with a 20 sec pause between each immersion. The wash steps are repeated in a second Wash plate before the beads are air-dried for 30 sec on the extractor before DNA/RNA is released into an Elution Plate (Eppendorf V-bottom microplate – E951040188) pre-filled with 50 μl of DEPC-treated water by dipping the extractor into the plate for 1 min. It is then removed and the cover plate is discarded at the plate-stripping station before the extractor is returned to its dock. If desired, larger lysate volumes (up to 5 ml) can be processed using a KingFisher Flex (Thermo Fisher).

Time to Complete: 1 h

Supplies: Lysis buffer, plates, SpeedBeads, primers

Cost: $0.41 USD per sample

Infrastructure: 1 Beckmann FX

#### 3. RT-PCR

RT-PCR targets a 177 bp segment of the E gene (Figure 3). A 96-well head on a FX liquid handler employs 20 μl tips (Beckman Coulter P20 AP96 barrier) to transfer 1 μl of RNA extract from four 96-well Elution plates into a 384-well plate (Eppendorff twin.tec LoBind skirted) prefilled with 5 μl of Superscript IV One-Step RT-PCR System (Thermo Fisher) spiked with 100 copies of a 368 bp gBlock synthetic DNA oligomer (Integrated DNA Technologies) which serves as an in-well positive control. The gBlock included two important modifications from the reference E gene sequence (26225..26493 GenBank NC_045512.2): a 100 bp random sequence was inserted 62 bp from the 5’ terminus and a short inversion was sited 111–132 bp from the 5’ terminus. Plates scheduled for Sanger sequencing also contain 0.1 μM of the F and R primers for the E gene (Integrated DNA Technologies) in each well. Plates destined for Sequel are pre-made in sets of 24 with each of the 9,216 wells containing a different primer pair (96F, 96R), each with a distinct 16 bp Unique Molecular Identifier (UMI), allowing every sequence to be linked to its source well (Figure 3). The same primer structure is used for the S5, but the UMI labelling is simpler because the wells in each 384-well plate can share the same set of primers (16F, 24R) because the amplicons from different plates are discriminated by secondary indexing after RT-PCR (Figure 2). Each 384-well plate includes four negative and four sample-free positive controls, the latter containing 100 copies of synthetic RNA for the E gene. After assembly, each plate is placed in a 384-well PCR block for RT-PCR (reverse transcription at 55°C for 10 min is followed by PCR cycling that begins with denaturation at 98°C for 2 min, followed by 50 cycles of 98°C for 10 sec, 55°C for 10 sec, and 70°C for 12 sec in the first 30 cycles versus 30 sec in the last 20 cycles, before a final extension at 72°C for 5 min). The extension time for the first 30 cycles of PCR is too brief to allow amplification of the gBlock, but the longer extension time allows its amplification during the final 20 cycles.

**Figure 3:**
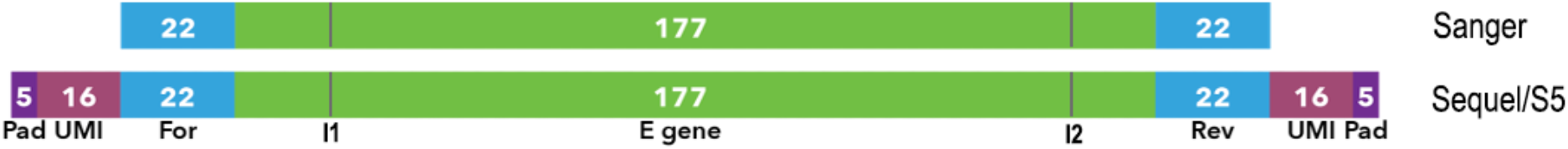
Diagram of the 177 bp segment of the E gene that is amplified using 22 bp Forward and Reverse primers in the RT-PCR for samples destined for Sanger sequencing. Samples scheduled for analysis on Sequel or S5 employ primers that include a 5 bp pad, a 16 bp Unique Molecular Identifier (UMI) as well as the E gene primers. The two lines indicate positions that were modified in the gBlock (I1 = 100 bp insertion, I2= 22 bp inversion).

Time to Complete: 1 h

Supplies: RT-PCR reagents, plates, tips, primers

Cost: $0.84 USD per sample

Infrastructure: 384-well PCR block for Sanger (16 for Sequel/S5); 1 Biomek FX

#### 4. E-Gel

Once RT-PCR is complete, the plate is returned to the FX deck where 3 μl is removed from each well and injected into a slot in four E-Gels (Thermo-Fisher). After being run for 6 minutes, the E-Gel is photographed (Figure 4) and the image is uploaded to the LIMS which assesses the presence/absence of amplicon(s) for each well. This is achieved by splitting the overall image into one image for each well. These individual records are standardized for variation in background by examining unoccupied segments of the gel before insert positions are inferred, band positions are assessed, and a band trace is constructed (Figure 4). This analysis detects the presence or absence of band (s) in each well trace and ascertains if they match the expected size of the E gene amplicon (221 bp for Sanger, 263 bp for Sequel and S5) and/or the gBlock control (368 bp) while also determining their intensities.

**Figure 4:**
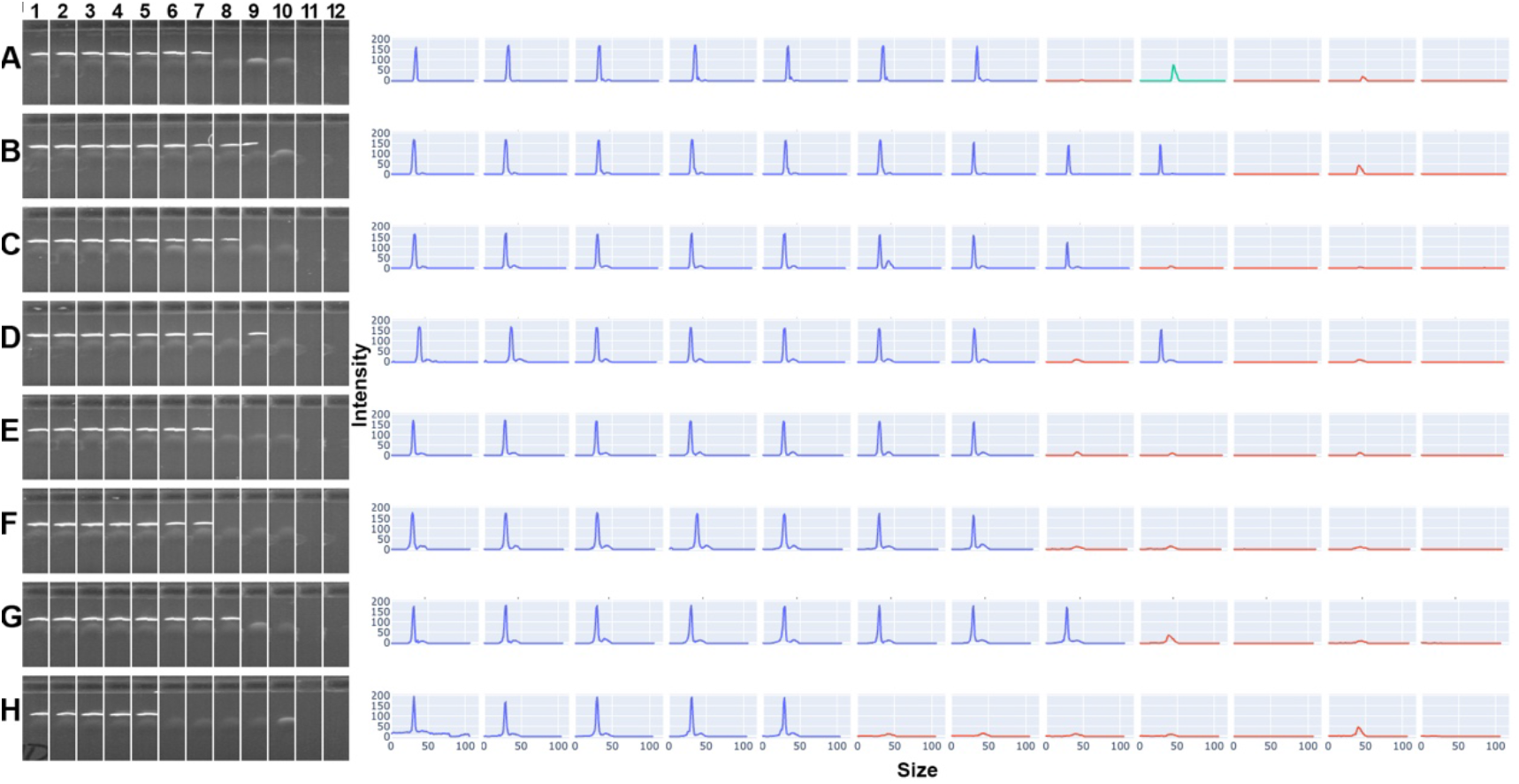
E-Gel showing recovery of amplicons for the E gene of COVID-19 on left and automated scoring of E-Gel for amplicon size and intensity on right. Intensity is maximal with bright white and zero for black. Because this analysis did not employ an in-well standard, positives show one band while negatives lack a band. Positive samples are A1-7, B1-8, C1-8, D1-7, D9, E1-7, F1-7, G1-8, and H 1-5.

##### Data Analysis

Four band configurations are possible for each E-Gel lane. Negative samples show a single band positioned at 368 bp derived from the gBlock control while positives can either possess two bands (one from the 221/263 bp amplicon of the E gene and one at 368 bp from the gBlock) or one band corresponding to the E gene when viral loads are high. Lanes lacking a band are false negatives that might reflect the presence of inhibitors in the RNA/DNA extract while bands with low intensity or the wrong size lead to an inconclusive assay.

##### Sensitivity

E-Gel screening was 100% successful in detecting positive controls with 50 or more RNA copies of the E gene. Success declined at lower concentrations (88% at 25 copies, 40% at 10 copies, 25% at 5 copies).

Time to Complete: 0.2 h

Supplies: E-Gels and tips

Cost: $0.30 USD per sample

Infrastructure: 4 E-Gel Stations, 1 Biomek FX

#### 5A. Sequencing Protocol 1 – Sanger

When sample throughput is less than 1,000 per day, Sanger sequencing allows cost-effective and rapid characterization of the amplicons generated by RT-PCR. This workflow involves two steps (sequencing reaction, data analysis).

##### Sequencing Reaction and Clean-up

Regardless of E-Gel results, the product in each well from RT-PCR is unidirectionally sequenced using BigDye v3.1 (Thermo Fisher 4337455) and the reverse E gene primer. Sequencing reactions are performed by adding 0.5 μl of each diluted RT-PCR reaction product (1:5 H_2_O to lower the high salt concentration) into 384-well plates prefilled with 5 μl of sequencing reaction mix (0.175 μl of BigDye, 1.2 μl of 5x sequencing buffer (400 mM Tris-HCl pH 9.0 + 10 mM MgCl_2_), 3 μl of 10% trehalose, 0.5 μl of 10 μM reverse primer, 0.125 μl of molecular grade water). The thermocycling regime employs initial denaturation at 96°C for 2 min, followed by 30 cycles of 96°C for 30 sec, annealing at 55°C for 15 sec, and extension at 60°C for 4 min.

Cycle sequencing products are purified using an automated method (Elkin et al. 2002) on a Biomek FX using SpeedBeads (GE healthcare 45152105050250) in TEG (Tetraethylene glycol, Sigma). 2 μl of magnetic bead suspension (7.5 mg/ml in 50% TEG) and 10 μl of 85% ethanol are added to each 6 μl cycle sequencing reaction and mixed 15 times by pipetting. The plate is incubated for 5 min at 20°C, transferred to a magnet, and incubated for 3 min before the supernatant is removed. The beads are washed twice with 30 μl of 85% ethanol and then dried for 10 min before the cycle sequencing products are eluted in 35 μl of 5 mM DTT and 0.5 mM NaHCO_3_. Finally, 15 μl from each well is transferred to a 384-well optical plate (Applied Biosystems) and loaded for analysis on an ABI 3730XL using a 1 hour run (IP300). Trace files are submitted to BOLD www.boldsystems.org) for interpretation.

##### Data Analysis

When a well generates a trace file, it is converted to a Phred file and assembled with CAP3 (Huang and Madan 1999) to automatically produce a sequence. Primers are trimmed and the sequence is uploaded to BOLD after quality filtering. Sequences are aligned to a reference sequence for the E gene and the 22 nucleotide positions (111–132 bp from 5’ start) corresponding to the inversion in the gBlock are examined to assess each sequence for heteroplasmy at these positions, a result indicating that it derives from the gBlock and E gene. Alternatively, a clean sequence for the gBlock control indicates the sample contained no E gene template while a clean sequence for the E gene indicates that high concentrations of viral RNA led to its dominance in the amplicon pool. In the first and third cases, the sequence for the E gene is extracted and queried against a reference library composed of all unique sequences of the E gene extracted from GenBank for SARS, MERS, and COVID-19. Any sample with a sequence with >99% similarity to >150 bp of the reference sequence for COVID-19 is designated as positive.

##### Sensitivity

Sanger analysis recovered sequences from 100% of positive controls with 10 or more copies of the E gene.

Time to Complete: 4 h

Supplies: Big Dye, SpeedBeads, plates

Cost: $1.64 USD per sample

Batch Size: 352

Infrastructure: ABI 3730XL Sequencer, Biomek FX, BOLD Platform

#### 5B. Sequencing Protocol 2 – Sequel

A Sequel run of multiplexed samples typically analyzes amplicons from 24 plates (9,216 samples) following a protocol for targeted amplicon sequencing (Hebert et al. 2018). This allows the screening of 8,832 samples and 384 controls (8 positives + 8 negatives per plate). The workflow involves two steps – preparation of circularized amplicons for SMRT sequencing and data analysis.

##### SMRT Sequencing

Amplicons from the 9,216 reactions are pooled before SMRTbell library preparation and the quality of the pool is assessed (Qubit, Bioanlayzer) prior to DNA damage- and end-repair. Repaired DNA is purified using AMPure beads before blunt end ligation. SMRTbell template prep kit v1.0 is used to generate circular sequencing templates and Binding Kit v3.0 is employed to attach the sequencing polymerase to primer-annealed SMRTbells. The resulting library is loaded on a SMRT Cell and run on Sequel or Sequel II using Sequencing Kit v3.0 for 4 hours.

##### Data Analysis

Each SMRT Cell generates about 0.2 million circular consensus sequences on Sequel and 1.6 million on Sequel II. This means an average coverage of 20x per well on Sequel or 150x on Sequel II if all 9,216 wells contain amplicons, but the read count is reduced by filtering. Sequences are analyzed using a cloud-based informatics pipeline supported by two platforms, mBRAVE (mbrave.net) and BOLD (boldsystems.org). Raw data are analyzed to generate circular consensus sequences which are assigned to their source well by examining their UMIs before the primers and UMIs are trimmed. They are then filtered for quality and length before being dereplicated. Wells with sequences from the gBlock control are enumerated to verify the effectiveness of the in-well controls. In addition, all presumptive E gene sequences from a well are compared against a reference library for the E gene from human Betacoronavirus taxa (DS-HBCORONA – doi.org/10.5883/DSHBCORONA). Sequences with >99% similarity to COVID-19 are assigned to it and the number of reads for the E gene from each well is recorded. Although negative controls and wells from samples lacking viral RNA should have zero reads, aerosols can introduce a few E gene templates into wells that would otherwise lack them so those producing very low counts (< 5) are classed as negatives.

##### Sensitivity

The stability of read counts and the incidence of false positives and negatives were assessed by running two sets of 384-well plates with layouts and analytical constraints designed to stringently test the risk of misclassification. The risk of false positives was maximized by intermingling positive and negative controls by spiking odd-numbered wells with the E gene while even-numbered wells were left blank. In addition, PCR conditions were varied among plates to produce variation in read counts. The latter approach led to 10-fold variation in read counts (3,199–32,893) among the 48 plates with the mean read count per positive well ranging from 17–172. When the amplicon pool from one set of 24 plates (n = 9,216) was analyzed on two SMRT Cells, the read counts for individual wells showed strong concordance between runs (Figure 5). The incidence of false positives and negatives was then investigated by examining read counts from two separately constructed amplicon pools (each = 9,216 wells) analyzed on two SMRT Cells. To simplify analysis, the two read counts from each well were combined. As expected, most of the 310,532 reads (99.76%) derived from wells with template, but 743 were from blank wells after filtering wells with <5 reads. Histograms of the read counts for all positive and negative wells showed limited overlap with a threshold of 5 reads effectively discriminating the two categories. A heat map (Figure 6) closely approximated the expected checkerboard pattern with 44 exceptions reflecting false negatives, all from plates with <10,000 reads (Table 1). The remaining exceptions involved false positives which averaged 3% in plates with <15,000 reads, but just 0.5% in those with more reads.

**Table 1:** Relationship between the incidence of false positives and false negatives for the E gene and the number of reads for each 384-well plate following analysis on Sequel. Positive status was assigned to all wells with five or more reads, negative status to those with fewer reads.

| Reads (K) | # Plates | # Wells | False Positives |  | % False Negatives |  |
| --- | --- | --- | --- | --- | --- | --- |
|  |  |  | Number | % | Number | % |
| <5 | 5 | 1920 | 15 | 1.56 | 32 | 3.33 |
| 5 – 10 | 7 | 2688 | 37 | 2.75 | 12 | 0.89 |
| 10 – 15 | 6 | 2304 | 50 | 4.34 | 0 | 0 |
| 15 – 20 | 2 | 768 | 2 | 0.52 | 0 | 0 |
| >20 | 4 | 1536 | 4 | 0.52 | 0 | 0 |

**Figure 5:**
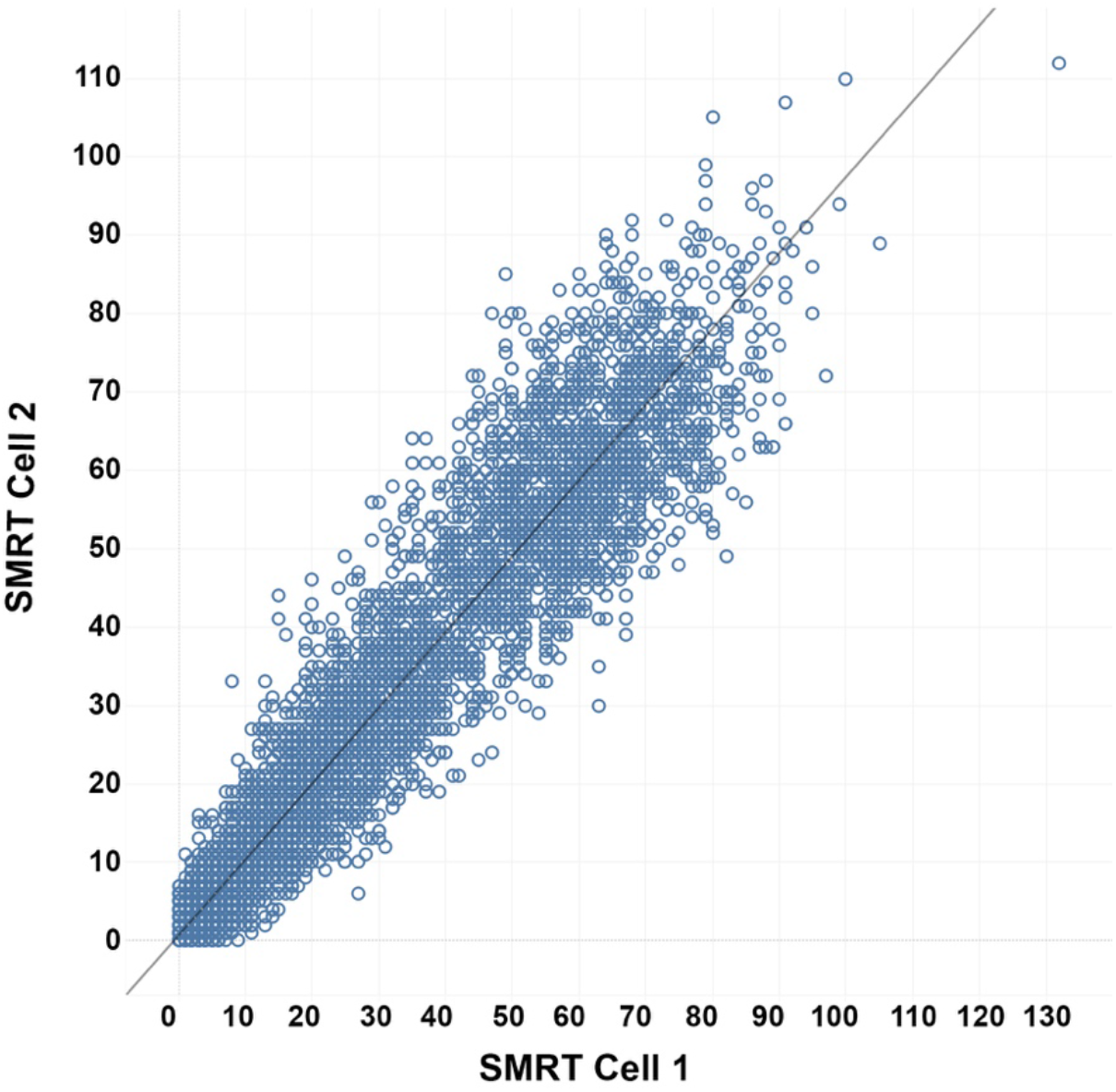
Correspondence in the number of read counts for the E gene from each of 9,216 wells when an amplicon pool was analyzed on two SMRT Cells.

**Figure 6:**
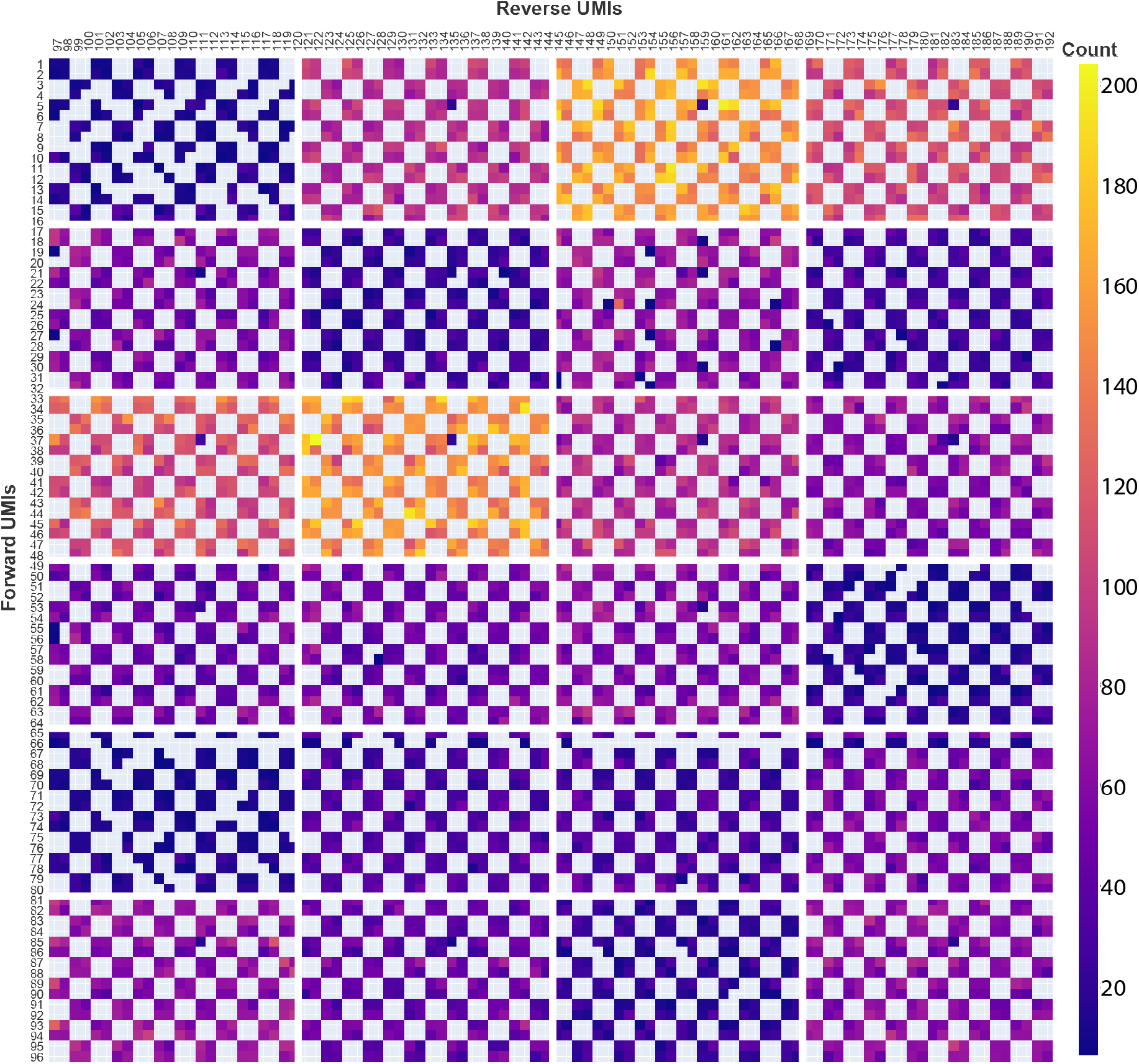
Heat map based on the total number of reads for the E gene recovered from each of 9,216 wells after analysis on two SMRT Cells. To aid visualization, each block represents the corresponding well in the four 96-well plates that were used to assemble each 384-well plate. The checkerboard pattern reflects the fact that alternate wells were either positive or negative controls for COVID-19.

Time to Complete: 36 h

Supplies: SMRT Cell and reagents

Cost: $0.22 USD per sample

Batch Size: 8,448

Infrastructure: 1 Sequel or Sequel II platform; BOLD and mBRAVE Platforms

#### 5C. Sequencing Protocol 3 – ION S5

Because of its high sequence output (10 million reads with 530 chip and ExT chemistry) and because Index ligation follows RT-PCR, individual runs of the S5 can analyze from 5–50 plates.

##### S5 Sequencing

The S5 workflow includes four steps: purification and quantification of the amplicon pool from each plate, index ligation, template preparation, and sequencing. The amplicons from each plate are tagged with a unique Index via the ligation of pool-specific Y-adapters (Forth and Hoper 2019) which also contain the necessary elements for sequencing. Following analysis, the E gene sequences are mapped to their source plate by examining their Index and to their source well by examining their UMIs. Because only the duration of the first two steps increases as number of plates rises, the overall analytical time is 19 hours for five plates versus 22 hours for 50.

##### S5 Data Analysis

Raw data are demultiplexed via their plate-level indices by the onboard S5 Torrent Browser before sequences are analyzed using a cloud-based informatics pipeline supported by two platforms, mBRAVE (mbrave.net) and BOLD (boldsystems.org). Each read is assigned to its source well by examining its two UMIs before the primers, adapters, and UMIs are trimmed. Sequences are then filtered for quality and length and dereplicated. Sequences that derive from the gBlock are enumerated as a test of the efficacy of the in-well standard. Presumptive E gene sequences are compared against a reference library for the E gene from human Betacoronavirus taxa (BOLD dataset – doi.org/10.5883/DSHBCORONA). Sequences with >98% similarity to COVID-19 are assigned to this taxon and the number of reads for the E gene from each well is enumerated.

Time to Complete: 20 h to 22 h

Supplies: Y-adapters (IDT), NEB Next Ultra II End Repair/dA-Tailing Module, ION 510, 520, 530 Sequencing Kit, 530 chip

Cost: $0.70 – $0.17 USD per sample

Batch Size: 1,760 to 17,600

Infrastructure: 1 S5 or S5 XL, ION Chef

##### Sensitivity

The incidence of misclassification was tested with a protocol similar to that for Sequel. The risk of false positives was maximized by spiking odd-numbered wells with the E gene while even-numbered wells were blank controls. In addition, varied ligation conditions produced variation in read counts among plates. In total, 2.38 million reads were recovered with a mean read count per plate of 99,362. Because there was 92-fold variation (2,559–239,888) in read count among plates (0–10K: 1; 10–20K: 3; 20–40K: 5; 40–80K: 3; 80–160K: 6; 160–320K: 6), the average number of reads per well ranged from 7–625 among the plates. As expected, most reads (98.22%) derived from wells with template, but 42,456 were from blanks. Because the plate with the lowest count lacked sequences from many positive wells, it was excluded from further analysis. Heat maps for the other plates closely approximated the expected checkerboard pattern (Figure 7). Among the 8,832 wells in these 23 plates, there were two false positives and 33 false negatives, an error rate of 0.4%. However, E-Gel results indicated that both false positives lacked a band while 32 of the 33 false negatives possessed one (Figure 8). Given this conflict between the two analytical endpoints, these 35 samples would be scored as ‘inconclusive’.

**Figure 7:**
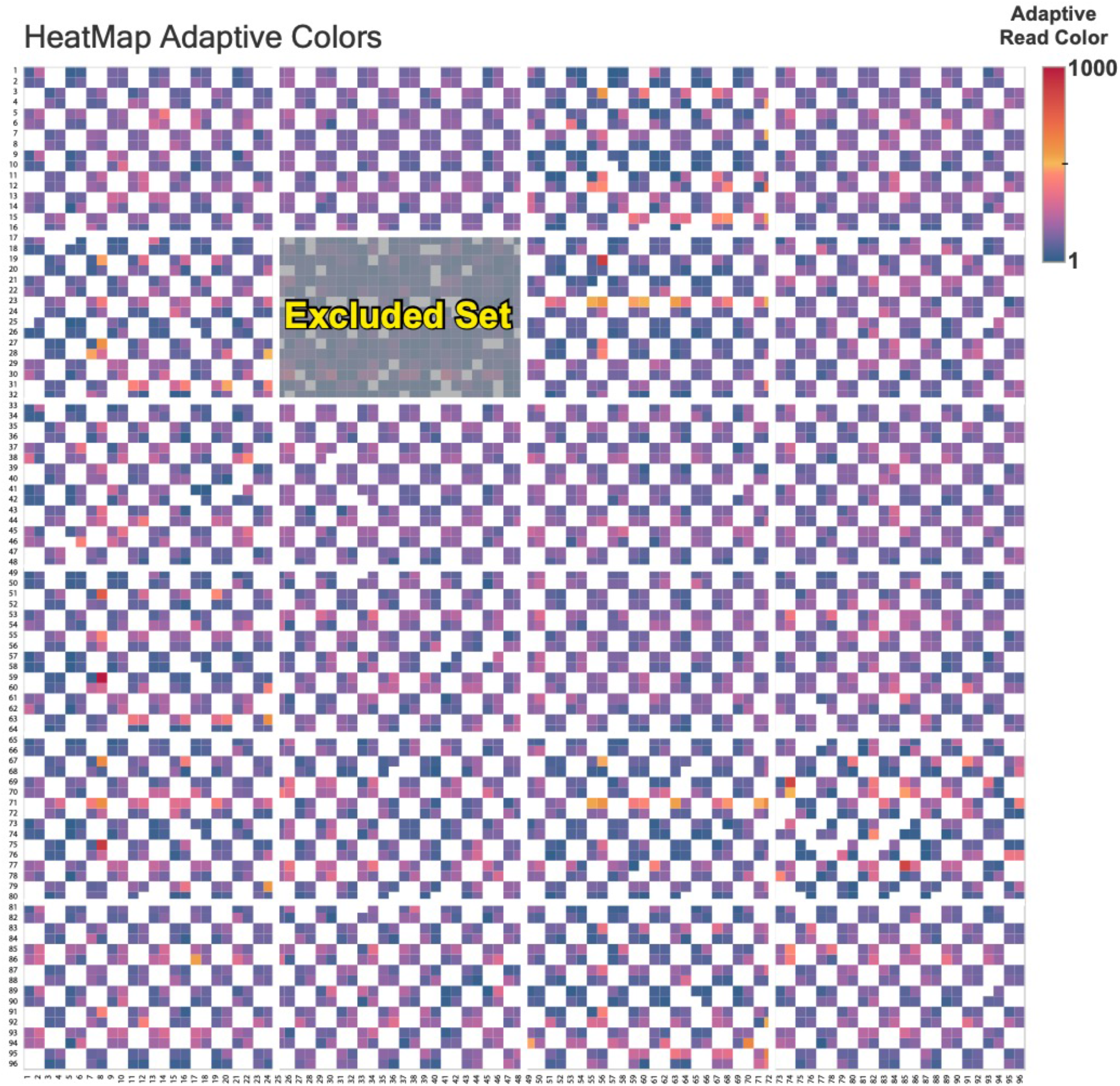
Heat map based on the total number of reads for the E gene recovered from each of 9,216 wells after analysis on the S5 (530 chip). To aid visualization, each block represents the corresponding well in the four 96-well plates that were used to assemble each 384-well plate. The checkerboard pattern reflects the fact that alternate wells were either positive or negative controls for COVID-19. The plate with the lowest count was excluded from analysis.

**Figure 8:**
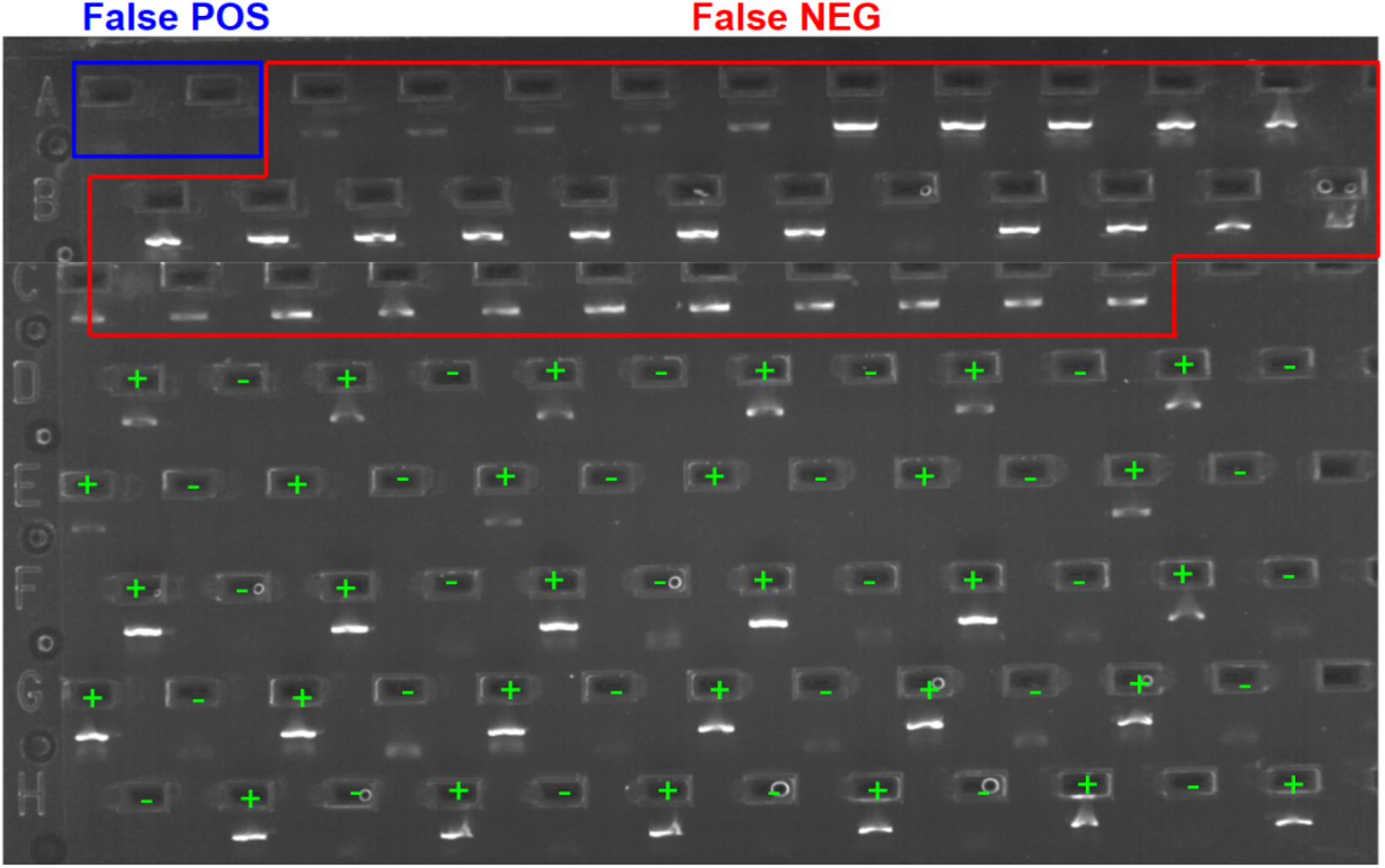
E-Gel showing results from the two false positives and 33 false negatives in the S5 run as well as 30 positive wells and 30 negative wells. The 30 positive wells were selected across the full range of sequence reads on the S5. Three of those with the lowest sequence counts failed to show a band on the E-Gel indicating greater sensitivity of the sequencing endpoint.

#### 6. Report Generation

Two reports are automatically generated for each sample; the first provides the E-Gel result (Figure 9) while the second provides the sequence result. Each report is generated as soon as the necessary data is available to produce them, allowing an initial diagnosis based on the E-gel, followed by verification through sequencing. Both reports include the sample code, its chain of custody including laboratory provenance, as well as a diagnosis, confidence interval, and results from the positive/negative controls in the run. Aside from these reports on individual samples, a summary report (Figure 10) is generated for each 384-well plate which provides diagnostic outcomes along with confidence intervals and results from positive/negative controls.

**Figure 9:**
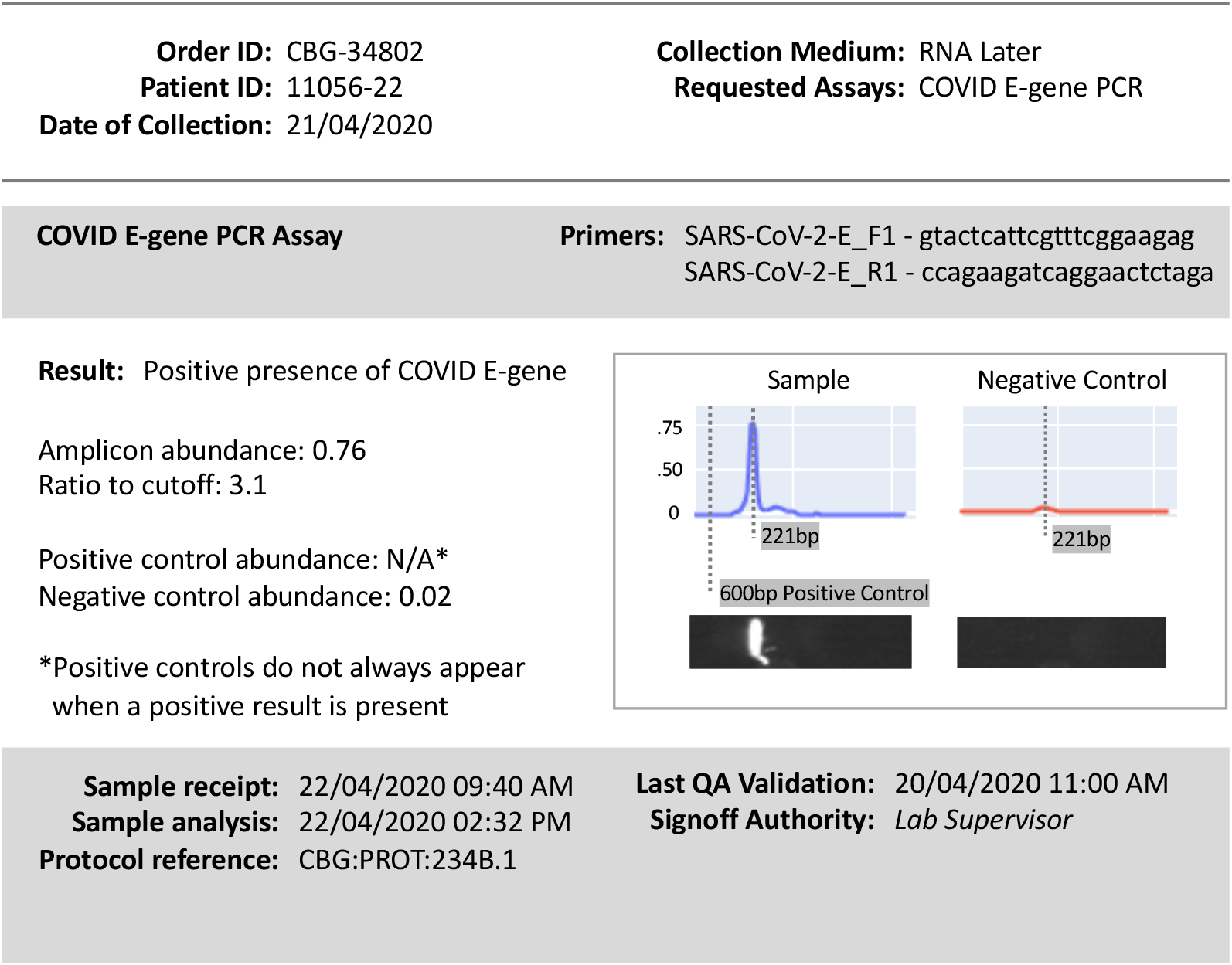
Sample report for an E-Gel assay.

**Figure 10:**
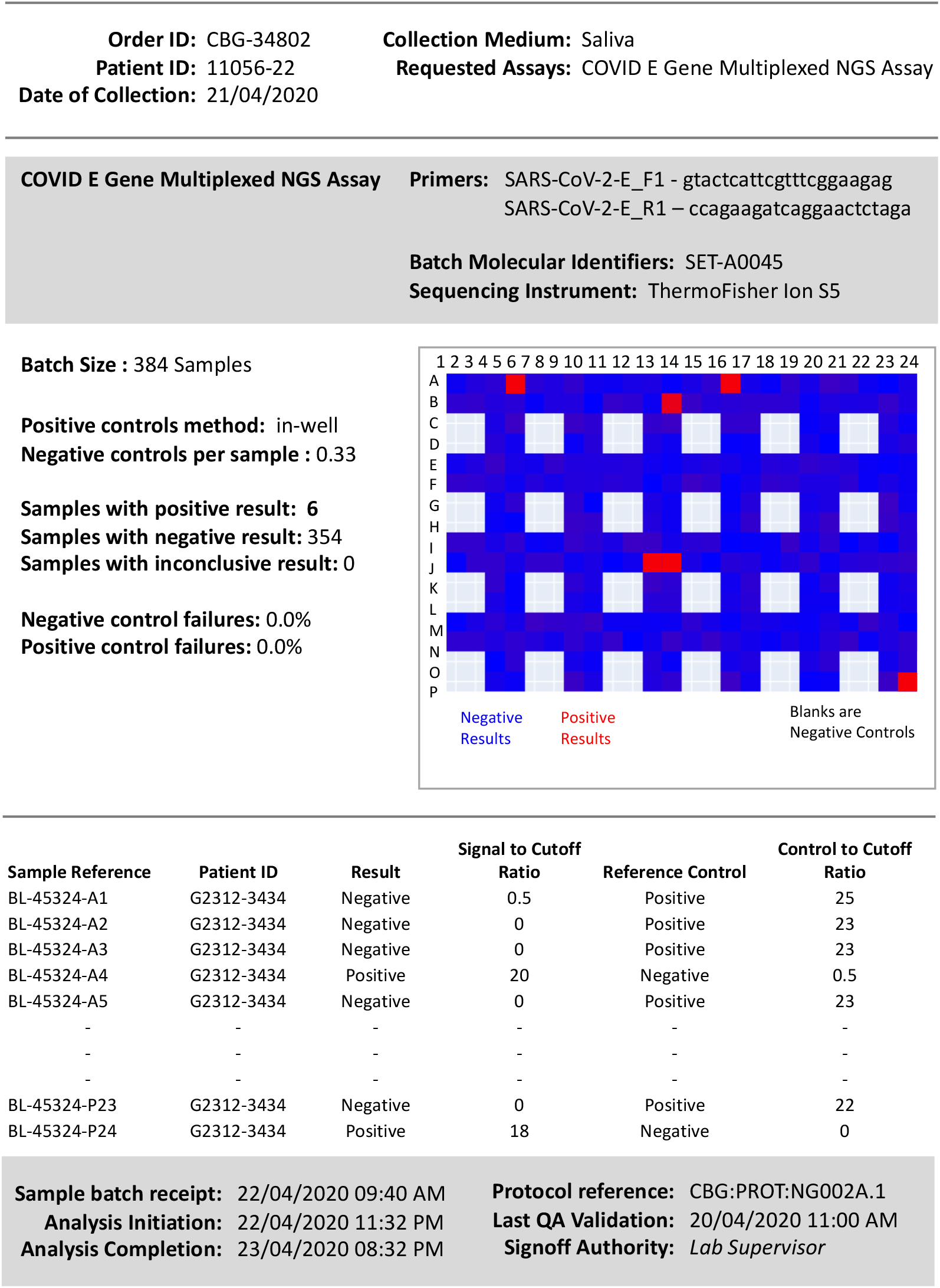
Batch report for a 384-well plate which combines results for E-Gels and sequencing.

## DISCUSSION

The devastating health impacts of COVID-19 have provoked efforts to slow its transmission by coupling testing programs with contact tracing. Where implemented early and pursued vigorously, these actions suppressed infection rates. However, limited analytical capacity and high costs have meant that only symptomatic individuals were tested in many nations, a strategy that has seen infection rates rise, in part because many individuals with COVID-19 are asymptomatic (Nishiura et al. 2020). In response, some nations (e.g. UK) have now adopted stringent physical distancing with the goal of reducing infections to the level where it becomes feasible to reactivate contact tracing and testing. An alternate solution involves immediate expansion of these activities. For example, the Rockefeller Foundation (2020) proposes the USA recruit 100,000 contact tracers and expand testing 30-fold within six months. With a weekly target of 30 million tests at $100 each, analytical costs alone are estimated at $150 billion annually. Given this projection, the need for less expensive tests is obvious so long as they also bring speed and reliability.

The current protocol employs a simple workflow to deliver a preliminary diagnosis from E-Gels within 8 hours of sample reception followed by a final decision based on sequence results within 24–48 hours. Fifty thousand samples can be processed in a standard work week, and production can be doubled with 16/7 operations without encountering an infrastructure barrier. By avoiding RNA extraction kits, employing low volume RT-PCR, and using high-throughput platforms for sequence characterization, both end points (Gel + sequence) can be achieved for less than $5 USD per sample. Moreover, in-well controls provide a robust measure of false negatives at both the E-Gel and sequence stage. Although both HTS platforms performed well, the S5 was more flexible because the indexing step provided a simple path to deal with day-to-day variation in the number of samples for analysis.

The protocols in this paper represent an important step towards a viral test that can be implemented for $1. In fact, if just one end point is pursued (E-Gel or sequencing), the material costs for the current protocol nearly meet this target ($1.55 for E-Gel, $1.42 for S5). Group analysis (Dorfman 1943, Edouard et al. 2015) can further reduce costs by 80% in settings where comprehensive testing is undertaken if infection rates are low (<2%). With this approach, pooled samples from multiple individuals are initially analyzed and individual testing only examines members of infected pools. By coupling the targeted analysis of a single gene region with group tests, comprehensive screening programs can be established at very low cost. This creates a new imperative; it is essential to identify inexpensive methods for sample acquisition as they currently exceed $10. The shift from nasopharyngeal to saliva samples is a key advance, but further work is needed to optimize protocols. In short, mass screening programs need innovation along the entire analytical chain from sample acquisition to report synthesis.

The control of COVID-19 or any similar pathogen relies upon suppressing its R_o_ below 1. Large-scale randomized testing programs are essential to monitor progress toward this goal, an undertaking which would be facilitated by the present approach. There is also a need for comprehensive testing in settings where confinement (e.g. old age homes, prisons) raises the risk of outbreaks. When it comes to broader societal monitoring of R_o_, schools provide an ideal venue because students represent 10-15% of the overall population and provide a window into the incidence of infections in a larger subsample (i.e. their parents). Moreover, their organization in groups of about 20 is ideal for group testing. For example, with a population of 15 million, Ontario has 2 million students who are educated in 100,000 classrooms. Given this demographic, group testing could assay infections in every class for less than $1 million per week, about 0.1% of the annual education budget (FAO 2019). Perhaps COVID-19 will be suppressed through the rapid development of a vaccine, but a quick exit from the current pandemic should not be an excuse to delay development of the capabilities needed to deal with a future pathogen whose suppression requires deep surveillance to reduce its R_o_.

## Supporting information

Fastq files for Ion S5 analysis of 24 indexed plates

Fastq file for Sequel analysis Run 1

Fastq file for Sequel analysis Run 2

## ACKNOWLEDGEMENTS

The study was enabled, in part, by awards from the Canada Foundation for Innovation, the Canada Research Chairs program, and the Ontario Ministry of Research and Innovation to PDNH, and by a grant (Food From Thought) from the Canada First Research Excellence Fund to the University of Guelph. We are also grateful to Ann and Christopher Evans whose support facilitated this work in important ways. We thank Suz Bateson for aid with graphics, and the informatics and sequencing staff at the Centre for Biodiversity Genomics for aiding data acquisition and analysis.

